# The Role of Anterior Insular Parvalbumin GABAergic Interneurons in Impulsivity-Related and Alcohol-Seeking Behaviors

**DOI:** 10.64898/2025.12.15.694283

**Authors:** F. Bezerra, G.J.D. Fernandes, A. Anjos-Santos, C.A. Favoretto, B.T. Rodolpho, P. Palombo, V. Vialou, K.P. Abrahao, F.C. Cruz

**Author notes:** Correspondence: Fernando Bezerra Romualdo da Silva, Departamento de Farmacologia, Universidade Federal de São Paulo, Rua Botucatu, 862, Ed. Leal Prado, 1st floor, São Paulo, SP 04023-062, Brazil. Fábio Cardoso Cruz, Departamento de Farmacologia, Universidade Federal de São Paulo, Rua Botucatu, 862, Ed. Leal Prado, 1st floor, São Paulo, SP 04023-062, Brazil.

## Abstract

Alcohol use is a major global health concern, particularly during adolescence, a developmental period characterized by heightened impulsivity, a known risk factor for alcohol misuse. The insular cortex is critically involved in interoceptive processing and impulsivity regulation, and its activity is powerfully shaped by parvalbumin (PV)-expressing GABAergic interneurons. We hypothesized that developmental differences in PV interneuron activity within the insular cortex contribute to impulsivity-related behaviors and alcohol consumption. To investigate this, male C57BL/6 mice were assessed at two developmental stages: adolescence (postnatal day 28) and young adulthood (postnatal day 64). Impulsivity was measured using the Omission-Contingency Learning (OCL) task, while alcohol intake and preference were evaluated using a two-bottle choice paradigm. Adolescent mice displayed higher impulsivity and greater alcohol consumption than adults. Double immunofluorescence labeling (Fos+/NeuN+ and Fos+/PV+) showed that increased impulsivity was associated with reduced activation of PV interneurons in the insular cortex. Chemogenetic activation of PV interneurons in adult PV-Cre mice reduced impulsivity in the OCL task and decreased alcohol intake. These findings indicate that insular PV interneurons play a modulatory role in impulsive behavior and alcohol consumption, suggesting a potential therapeutic target for impulse-control and alcohol-use disorders.

## 1. Introduction

Alcohol use remains one of the most widespread forms of psychoactive substance misuse worldwide and continues to represent a critical public health challenge [1], [2]. Globally, approximately 3 million deaths, accounting for 5.3% of all annual mortality, are attributed to harmful alcohol consumption [3]. Alcohol use disorder (AUD) is highly prevalent and is defined by compulsive intake, diminished behavioral control, and persistent consumption despite adverse personal, social, and physiological consequences [4]–[6]. Importantly, the initiation of alcohol use typically emerges during adolescence, a developmental window marked by substantial neurobiological remodeling [7]. Adolescence is associated with heightened risk-taking, emotional reactivity, and increased impulsivity, characterized by a preference for immediate rewards and reduced inhibitory control [8]–[10]. Exposure to alcohol during this critical period may amplify these vulnerabilities, reinforcing maladaptive reward-driven behaviors and increasing susceptibility to the development of AUD [11], [12]. Finally, recent studies link impulsivity to the onset and escalation of alcohol use [13], with more impulsive adolescents showing greater tendencies toward hazardous drinking and excessive consumption later in life [14]–[16].

At the neural level, the insular cortex functions as a key integrative hub, synthesizing interoceptive, affective, cognitive, and executive signals. Its extensive anatomical and functional connections with frontal cortical regions and the basal ganglia support its essential role in both adaptive and maladaptive impulsive behaviors [17]–[19]. Clinical and preclinical studies have demonstrated that alterations in insular functional connectivity particularly with frontal and limbic structures are associated with impulsivity-related phenotypes[20], [21]. Developmental changes in the structural and functional maturation of the insular cortex during adolescence have also been linked to impulsivity, with variations in cortical thickness and neuronal excitability emerging as potential markers of vulnerability to compulsive behaviors. Recent findings indicate that the reduced excitability of anterior insula neurons during adolescence is correlated with increases in impulsive and compulsive behaviors [22]– [24]. In contrast, chemogenetic activation of this region attenuates such behavioral responses [25].

Within the insular cortex, excitatory pyramidal neurons and inhibitory GABAergic interneurons maintain the delicate balance between excitation and inhibition required for proper cortical processing[26], [27]. Parvalbumin-positive (PV) interneurons, which provide fast and potent perisomatic inhibition that regulates pyramidal neuron output and supports synchronous cortical activity [28]. This excitation– inhibition balance is essential for normal cognitive function and is disrupted across multiple neuropsychiatric conditions [29], [30]. Recent studies have demonstrated that chemogenetic manipulation of PV interneurons in regions such as the medial prefrontal cortex and anterior cingulate cortex can modulate reward-seeking behavior, compulsive behavior, attention, and impulsivity [31], [32]. In the insula specifically, alterations in PV interneuron density or myelination have been associated with impaired danger recognition and enhanced pursuit of immediate rewards [30]–[32]. However, despite these emerging observations, the contribution of insular PV interneurons to impulsive behavior remains insufficiently understood.

Therefore, the present study aims to investigate the role of PV interneurons in the insular cortex during adolescence, a critical developmental stage characterized by heightened impulsivity and increased vulnerability to alcohol use disorder.

## 3. Materials and Methods

### 3.1 Animals

Transgenic PV-Cre mice on a C57BL/6 genetic background, as previously described [33] were obtained either from the Center for Development of Experimental Models for Biology and Medicine (CEDEME, Universidade Federal de São Paulo, São Paulo – UNIFESP) or from The Jackson Laboratory (B6;129P2-Pvalb^tm1(cre)Arbr^/J). Both lines were backcrossed onto the C57BL/6J mice. Animals were tested at two developmental stages: adolescence, defined as the period surrounding sexual maturation (postnatal days 28–47), and young adulthood (approximately postnatal day 64) [34]. Mice were housed under a reverse 12–h/12–h light–dark cycle (lights off at 06:00) in a temperature- and humidity-controlled facility (21 ± 2 °C). During behavioral testing, animals were maintained at 85% of their baseline body weight through controlled food restriction. All experimental procedures followed the Principles of Care of Laboratory Animals and were approved by the Ethics Committee on Animal Use at UNIFESP (CEUA #9191091219) as well as by the European Committee Council Directive 2010/63/EU, with authorization from the Animal Experimentation Ethics Committee C2EA-05 Charles Darwin and the French Ministry of **Higher Education and Research.**

### 3.2 Omission-Contingency Learning (OCL) Test

The OCL task was performed in operant conditioning chambers (170 × 150 × 200 mm, Med Associates®, VT, USA) placed inside sound-attenuated boxes. Each chamber was equipped with a yellow cue light (7.5 W) positioned above the active nose-poke (ANP) aperture, a white house light (7.5 W; 12 lux), and a single ANP positioned 45 mm above the floor on the same wall as the reward magazine. The magazine was connected via polyethylene tubing to a pellet dispenser (ENV-203IR, Med Associates®). Chambers and dispensers were controlled by MED-PC IV software, which recorded operant responses and delivered both cue light signals (3 s) and sucrose pellets.

Animals were trained under schedules of increasing fixed-ratio (FR) FR1, FR3, and FR5. The test phase consisted of a single 30-minute session conducted 24 hours after the final FR5 session. During the test, the software was programmed to deliver one sucrose pellet every 30 s; however, any ANP response performed before pellet delivery reset the 30-s interval. The primary outcomes measured were the total number of ANP responses and the number of rewards (REW) earned.

### 3.3 Two-bottle choice and quinine adulterated drinking sessions

To evaluate alcohol consumption and preference, mice were individually housed in cages equipped with two 50-ml conical plastic bottles. Animals had simultaneous access to one bottle containing tap water and another containing a 10% (v/v) ethanol solution. Twelve daily drinking sessions were conducted, each lasting 4 h (08:30–12:30). During the first six sessions (S1–S6), mice were presented with water and 10% ethanol. Subsequently, animals underwent three quinine-adulteration sessions in which the ethanol bottle contained 10% ethanol mixed with increasing concentrations of quinine (0.6, 0.8, and 1.0 mM across consecutive sessions). After the quinine phase, mice completed three additional sessions with access to water and 10% ethanol. The variables measured were alcohol consumption (g / kg) and ethanol preference, calculated as: [(volume of ethanol consumed) / (volume of alcohol solution + volume of water consumed)] × 100.

### 3.4 Immunofluorescence

Coronal brain sections (30 µm) were collected on a cryostat and rinsed five times (5 min each) in 0.1 M phosphate buffer (PB). Antigen retrieval was performed by incubating the sections in 0.01 M sodium citrate buffer (pH 8.0) at 70 °C for 20 min, followed by a 10-minute cooldown at room temperature. Afterward, the slices were washed in PBS and incubated in 1% hydrogen peroxide for 20 minutes under gentle agitation. Sections were rewashed and incubated for 2 h with biotinylated goat anti-mouse IgG (1:1000; #BA9200, Vector Laboratories) prepared in PB containing 1% Tween-20 and 3% normal goat serum. Following additional PB washes (5 times, 5 min per wash), sections were incubated for 48 h at 4 °C with combinations of the following primary antibodies diluted in PB with 1% Tween-20 and 2% powdered milk: anti-Fos (1:3000; rabbit, Cell Signaling Technology; or mouse, Abcam), anti-NeuN (1:2000; mouse, Millipore), and anti-PV (1:3500; rabbit, Abcam).

After primary incubation, sections were washed and incubated for 2 hours at room temperature with Alexa Fluor 488 (1:200 dilution; anti-rabbit or anti-mouse; Invitrogen) or Alexa Fluor 594 (1:1000 dilution; anti-mouse or anti-rabbit; Invitrogen) secondary antibodies, protected from light. Slices were then rinsed and incubated with DAPI (1:10,000) for nuclear labeling. Following PB washes, sections were mounted on gelatin-subbed slides, allowed to dry, and then coverslipped with Fluoromount (Sigma-Aldrich). Slides were stored horizontally at 4–8 °C for 48 h before being sealed with nail polish.

Images were acquired using a Zeiss Axioskop 2 microscope equipped with an Axiocam 503 color camera. Percent neuronal activation was quantified using the formulas: [(overlap of cells positive for Fos and specific neuronal marker (NeuN) / number of cells positive for NeuN) * 100]. The calculation of the percentage of PV-activated cells was performed using the following formula: [(overlap of cells positive for Fos and PV+) / number of cells PV+) * 100].

### 3.5 Immunohistochemistry

Coronal brain slices (30 µm) were collected following stereotaxic coordinates from the Paxinos and Watson (2007) atlas. Sections containing the regions of interest were rinsed in 0.1 M PBS, incubated in 3% hydrogen peroxide for 20 min, and washed again. Slices were then incubated for 24 hours at 4°C in anti-Fos primary antibody (1:4000; rabbit; Cell Signaling Technology) diluted in blocking solution (0.25% Triton X-100 and 3% normal goat serum in PBS). After washing, sections were incubated for 2 hours at room temperature with biotinylated anti-rabbit secondary antibody (1:1000; Vector Laboratories). Following additional washes, slices were incubated for 90 min in an avidin-biotin peroxidase solution (ABC Elite Kit, Vector Laboratories). Fos immunoreactivity was visualized using 3,3′-diaminobenzidine (DAB) for ∼10 min. Sections were then washed, mounted on gelatin/chromic aluminum-subbed slides, dried, and dehydrated through ascending ethanol concentrations and xylol. Slides were coverslipped with Entellan (Merck).

Images were captured using a Zeiss Axioskop 2 bright-field microscope with a 10× objective. Fos-positive nuclei were quantified from eight hemispheres (four slices per mouse) using QuPath software. Image acquisition and quantification were performed in a blinded manner, independent of the experimental group.

### 3.6 Surgical procedures

Stereotaxic surgeries were performed on PV-Cre BAC transgenic mice (PND 35) under isoflurane anesthesia (4% induction, 1.5–3% maintenance), with subcutaneous lidocaine administered for local analgesia. Cre-dependent adeno-associated viral (AAV) vectors were obtained from Addgene (Watertown, MA, USA). Chemogenetic inhibition and excitation were achieved using AAV1-hSyn-DIO-hM4D(Gi)-mCherry (Addgene viral prep #44362-AAV1; RRID:Addgene_44362) and AAV1-hSyn-DIO-hM3D(Gq)-mCherry (Addgene viral prep #44361-AAV1; RRID:Addgene_44361), respectively. AAV1-EF1a-DIO-eYFP (Addgene viral prep #27056-AAV1; RRID:Addgene_27056) was used as a Cre-dependent fluorescent control vector. It was unilaterally injected (0.2 µl/hemisphere) into the anterior insular cortex (coordinates relative to bregma: AP +1.7 mm, ML ±2.1 mm, DV −3.3 mm). Postoperative care included intramuscular streptomycin/penicillin (560 mg/ml/kg) and subcutaneous flunixin meglumine (0.5 mg/ml/kg). Viral constructs driven by synapsin or EF1α promoters ensured neuron-specific expression, enabling chemogenetic manipulation via DREADDs. Histological verification confirmed viral spread and targeting accuracy. Injections were counterbalanced between left and right hemispheres to account for functional lateralization of the insular cortex [35], [36].

## 4. Experimental design

The aim of Experiment 1 was to assess the impulsivity-related behaviors in adolescent and adult C57BL/6J male mice and to examine their relationship with alcohol consumption and preference (Fig 1A). For this purpose, twenty-nine adolescents (PND 28 at the beginning of the experiment) and thirty adults (PND 64 at the beginning of the experiment) were submitted to the OCL task. Forty-eight hours later, the same animals underwent the two-bottle choice and quinine-adulterated drinking tests.

**Figure 1.**
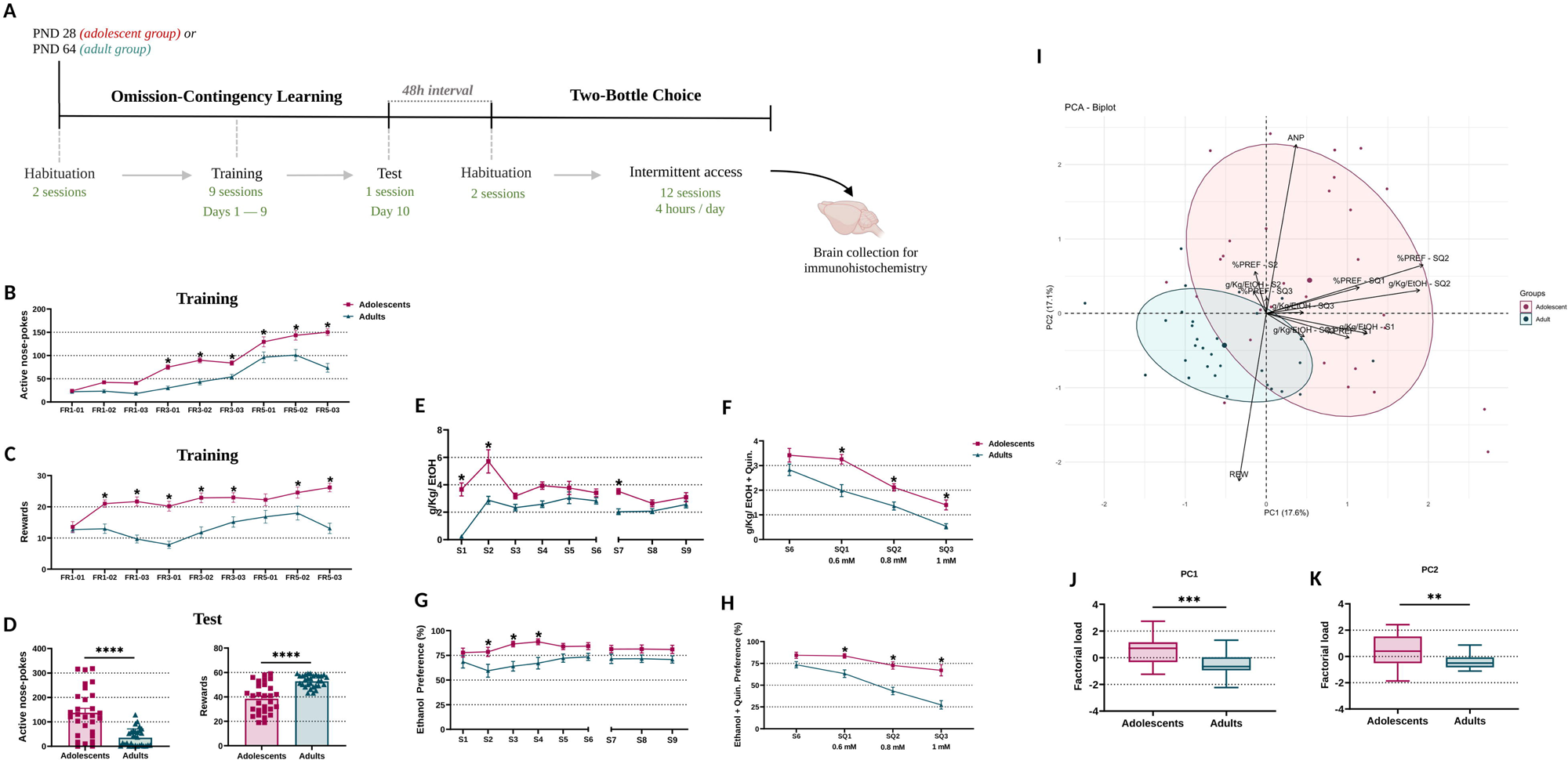
Omission-contingency learning and ethanol preference in adolescent and adult mice. (A) Schematic representation of the experimental design. Adolescent and adult mice underwent omission-contingency learning (OCL) training across nine consecutive sessions (Days 1–9), followed by behavioral testing on Day 10. Animals were subsequently evaluated in the two-bottle choice paradigm, including ethanol consumption, quinine-adulterated ethanol consumption, and ethanol preference assessments, as described in the Methods section. (B–C) Training phase of the OCL task. Adolescent mice displayed increased active nose-pokes (ANP) and rewards earned compared with adults across training sessions. (D) Test phase of the OCL task showing increased ANP and reduced rewards earned in adolescent mice, indicating impaired inhibitory control during omission-contingency testing. (E–F) Ethanol intake (g/kg) during baseline ethanol sessions (S1–S6 and S7–S9) and during quinine-adulterated ethanol sessions (SQ1–SQ3). Quinine concentrations are indicated on the x-axis. (G–H) Ethanol preference (%) during baseline ethanol sessions and quinine-adulterated ethanol preference (%) during SQ1–SQ3 sessions. (I) Principal component analysis (PCA) showing segregation of behavioral variables between adolescent and adult groups. (J–K) Variable loadings for PC1 and PC2, demonstrating stronger associations between ANP and rewards earned in adolescent mice compared with adults. Data are presented as mean ± SEM. Adolescents: n = 29; adults: n = 30. Statistical analyses were performed using two-way ANOVA or one-way ANOVA followed by appropriate post hoc tests. *p < 0.05, **p < 0.01, ***p < 0.001.

Experiment 2 was designed to assess the activation of PV GABAergic interneurons following the OCL task (Fig. 2A). A separate cohort of twenty-nine adolescents (PND 28) and thirty adults (PND 64) completed the OCL test and were perfused 1 h afterward. Double-label immunofluorescence was performed using Fos (a marker of neuronal activation) and NeuN (a marker of neuronal nuclei) to quantify neuronal activation, and Fos and PV to identify activated PV interneurons.

**Figure 2.**
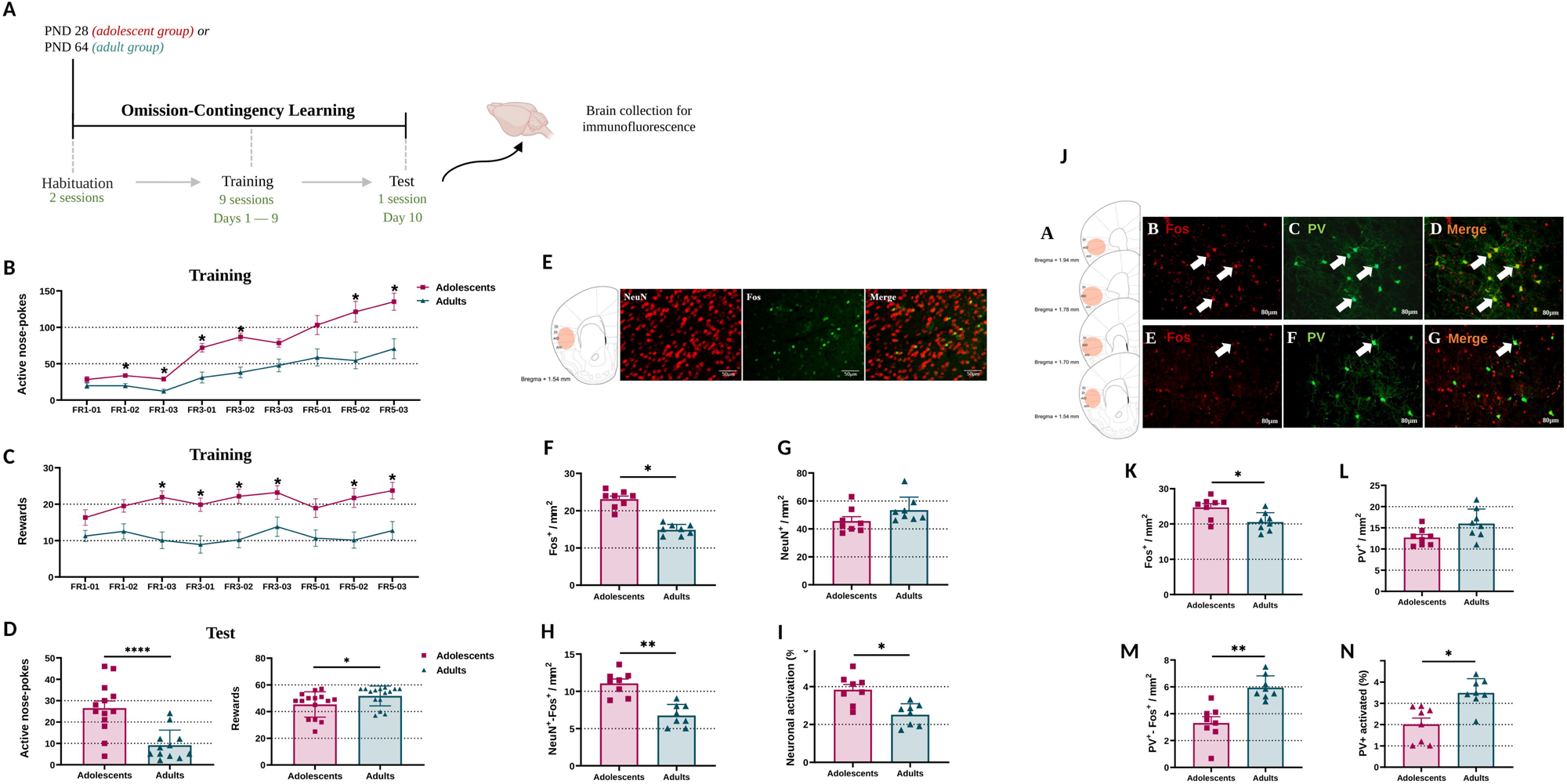
Insular cortex neuronal and parvalbumin interneuron activation following omission-contingency learning. (A) Experimental design for omission-contingency learning (OCL) followed by immunohistochemical and immunofluorescence analyses of the insular cortex. Adolescent and adult mice were trained and tested in the OCL task, and brain tissue was collected 1 h after testing. (B–C) During OCL training, adolescent mice exhibited increased active nose-pokes (ANP) and rewards earned compared with adults across sessions. (D) During the OCL test session, adolescent mice displayed increased ANP and reduced rewards earned relative to adults. (E) Representative confocal images of insular cortex sections showing NeuN-positive neurons (red), Fos-positive cells (green), and merged Fos+/NeuN+ co-localization (yellow). Scale bar = 80 μm. (F–I) Quantification of neuronal activation in the insular cortex. (F) Number of Fos-positive cells. (G) Number of NeuN-positive cells. (H) Number of Fos+/NeuN+ double-labeled cells. (I) Percentage of activated neurons (Fos+/NeuN+ relative to total NeuN+ cells). (J) Atlas representation of the analyzed insular cortex subregions (orange). Representative immunofluorescence images from adolescent and adult mice showing DAPI staining (blue), parvalbumin-positive (PV+) interneurons (green), Fos-positive cells (red), and merged Fos+/PV+ co-localization. Arrows indicate double-labeled cells. Scale bar = 80 μm. (K–N) Quantification of PV interneuron activation in the insular cortex. (K) Number of Fos-positive cells. (L) Number of PV+ cells. (M) Number of Fos+/PV+ double-labeled cells. (N) Percentage of activated PV interneurons. Data are presented as mean ± SEM. Behavioral analyses: n = 15 per group. Immunohistochemical analyses: n = 8 per group. Statistical analyses were performed using two-way ANOVA or one-way ANOVA followed by appropriate post hoc tests. *p < 0.05, **p < 0.01, ***p < 0.005, ****p < 0.0001.

Experiment 3 investigated the functional role of insular PV interneurons in impulsive behavior through chemogenetic manipulation (Fig. 3A). Thirty-nine PV-Cre mice received stereotaxic injections of viral vectors AAV1-hSyn-DIO-hM3DGq-mCherry, n = 13; AAV1-hSyn-DIO-hM4DGi-mCherry, n = 13; AAV1-EF1a-DIO-eYFP, n = 13) at PND 35. After a 21-day recovery period, mice completed the OCL task following intraperitoneal administration of CNO (1 mg/kg) 30 minutes before testing. A subset of animals (n = 8 / group) then underwent four forced-alcohol-consumption sessions (12, 8, 6, and 4 h) before repeating the OCL task, in which each nose-poke delivered 0.014 ml of 10% ethanol. Mice again received CNO before testing. Ninety minutes after testing, the mice were perfused, and their brains were collected for immunofluorescence analysis (Fig. 4A).

**Figure 3.**
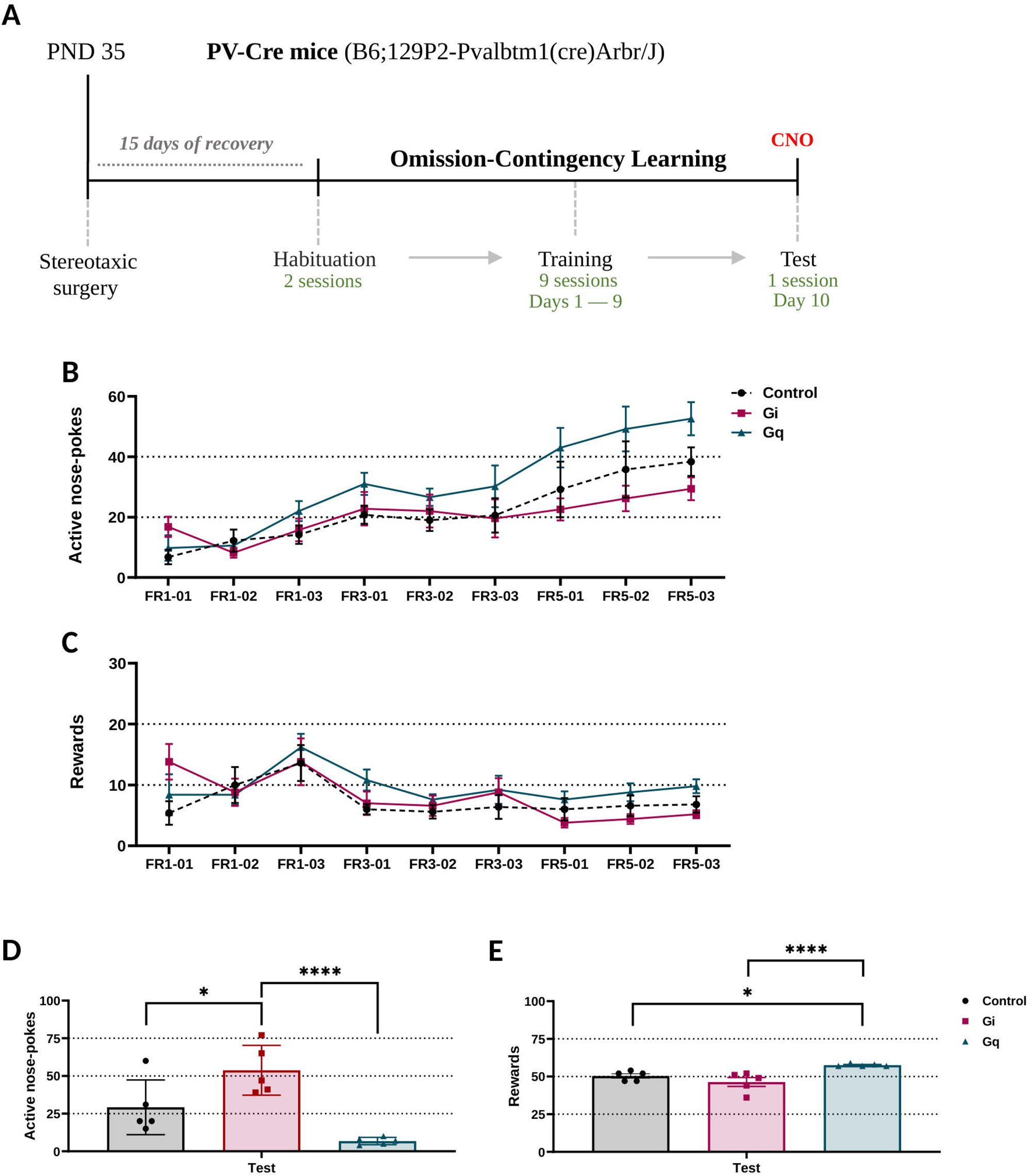
Chemogenetic modulation of insular cortex PV interneurons during omission-contingency learning. (A) Schematic representation of the experimental design for chemogenetic manipulation during omission-contingency learning. PV-Cre mice (B6;129P2-Pvalbtm1(cre)Arbr/J) received stereotaxic injections of Cre-dependent DREADD vectors targeting the anterior insular cortex, followed by a recovery period and behavioral testing under clozapine-N-oxide (CNO) administration. The timeline indicates training sessions across Days 1–9 and behavioral testing on Day 10. (B–C) Training phase of the OCL task showing active nose-pokes (ANP) and rewards earned across sessions in control, inhibitory DREADD (Gi), and excitatory DREADD (Gq) groups. (D–E) Test phase of the OCL task. Chemogenetic inhibition of PV interneurons increased ANP and reduced rewards earned, whereas chemogenetic activation produced the opposite behavioral profile. Data are presented as mean ± SEM. Control DREADD: black; Gi DREADD: red; Gq DREADD: blue. n = 5 per group. Statistical analyses were performed using two-way ANOVA or one-way ANOVA followed by appropriate post hoc tests. *p < 0.05, **p < 0.01, ****p < 0.0001.

**Figure 4.**
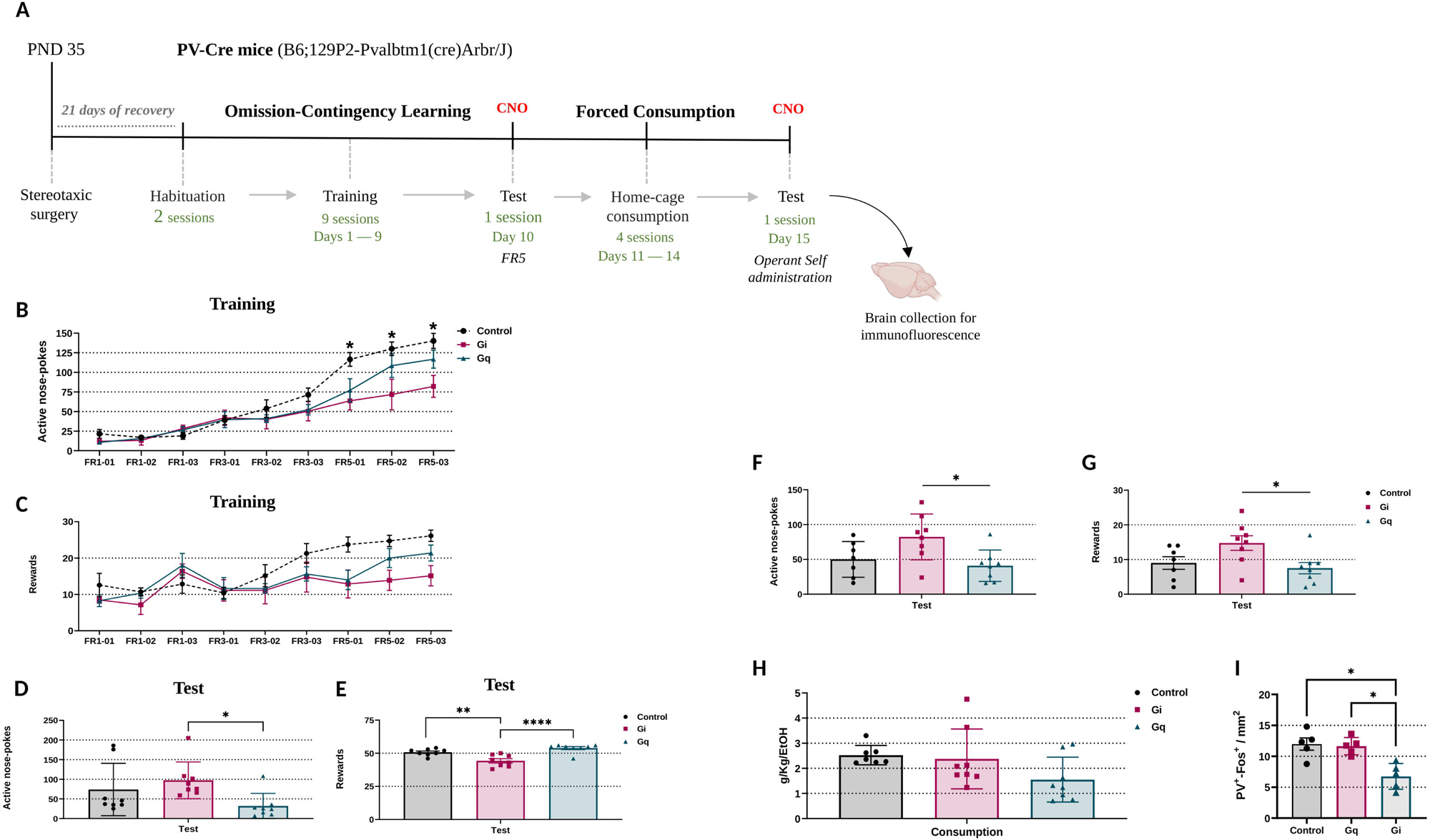
Chemogenetic modulation of insular cortex PV interneurons alters ethanol self-administration and neuronal activation. (A) Experimental design for chemogenetic manipulation during omission-contingency learning combined with ethanol intake procedures. PV-Cre mice received stereotaxic injections of Cre-dependent DREADD vectors into the anterior insular cortex, followed by recovery, CNO administration, operant training, forced ethanol consumption, and ethanol self-administration testing, as detailed in the Methods section. (B–C) During OCL training, mice expressing excitatory DREADDs (Gq) displayed increased active nose-pokes (ANP) and rewards earned compared with control and inhibitory DREADD (Gi) groups. (D–E) During the OCL test phase, chemogenetic inhibition of PV interneurons reduced ANP and rewards earned, whereas activation of PV interneurons increased behavioral responding. (F–G) Operant ethanol self-administration. Chemogenetic inhibition of PV interneurons reduced ethanol-seeking behavior and rewards earned compared with Gq mice. (H) Ethanol intake during the final habituation session showed no significant differences between groups. (I) Quantification of Fos+/PV+ double-labeled cells in the insular cortex following ethanol self-administration. Chemogenetic inhibition reduced PV interneuron activation, whereas chemogenetic activation increased Fos+/PV+ co-localization. Data are presented as mean ± SEM. Behavioral analyses: n = 8 per group. Immunohistochemical analyses: n = 5 per group. Statistical analyses were performed using two-way ANOVA or one-way ANOVA followed by appropriate post hoc tests. *p < 0.05, **p < 0.01, ***p < 0.001, ****p < 0.0001.

## 5. Data Analysis

All datasets were first assessed for normality using the D’Agostino–Pearson omnibus test and for homogeneity of variances using the F-test. Non-parametric comparisons were performed using the Mann–Whitney U test. For normally distributed datasets involving multiple factors or repeated measures, one-way or two-way repeated-measures ANOVA followed by Bonferroni *post-hoc* tests was applied. Animals were excluded (n = 7) only when identified as statistical outliers based on Grubbs’ test. All results are presented as mean ± standard error of the mean (SEM), and statistical significance was set at *p* < 0.05, *p* < 0.01, or *p* < 0.001.

For Experiment 1, training performance across sessions was analyzed using a two-way ANOVA (ontogenic period × session) with Bonferroni *post-hoc* adjustments. OCL test outcomes were evaluated using the Mann–Whitney U test. Prior to performing the principal component analysis (PCA), a Pearson correlation matrix was constructed using all OCL and two-bottle choice variables. Variables exhibiting high correlations (|*r*| > 0.8) were excluded to reduce redundancy. The PCA was then conducted using the remaining variables, and components were retained based on eigenvalues > 1.

For Experiment 2, the training data were analyzed using a two-way ANOVA (ontogenic period × session), followed by Bonferroni *post-hoc* tests. OCL test measures and neuronal activation data were analyzed using the Mann–Whitney U test.

For Experiment 3, OCL training data were analyzed using a two-way ANOVA (DREADD group × session), followed by Bonferroni *post-hoc* comparisons. OCL test outcomes, ethanol-seeking behavior, and locomotor activity were analyzed using one-way ANOVA followed by Tukey’s *post-hoc* test.

All statistical analyses were performed using SPSS (version 25.0.0) and Jamovi (version 1.2.27). Graphs were generated using GraphPad Prism 8 (version 8.0.2). The significance threshold for all analyses was α = 0.05.

## 4. Results

### 4.1 Adolescent mice exhibit greater impulsivity-related behaviors and ethanol consumption than adults

To determine whether adolescent mice exhibit heightened impulsivity and increased ethanol consumption compared to adults, we first analyzed their performance in the Omission-Contingency Learning (OCL) training sessions. For active nose-pokes (ANP; Fig. 1B), two-way ANOVA revealed significant differences for the ontogenic period factor [F(8,456) = 102; p < 0.0001] and interaction between factors [F(8,456) = 7.678; p < 0.0001. *Post-hoc* comparisons revealed that adolescents made significantly more ANPs than adults across all FR3 sessions (*p* < 0.05) and all FR5 sessions (p < 0.01), but not during FR1 training (p < 0.05). For the number of rewards (Fig. 1C), ANOVA likewise showed significant effects for the ontogenic period [F(8,456) = 11.94; *p* < 0.0001] and for the interaction between factors [F(8,456) = 6.322; *p* < 0.05], with group differences on the remaining training days (*p* < 0.05), with adolescents earning more rewards on all FR3 and FR5 training days except the first session.

In the OCL test session, Mann–Whitney analyses demonstrated robust age-related differences in both ANP and REW measures (*p* < 0.0001 for both), indicating impaired operant inhibition in adolescents compared to adults (Fig. 1D).

During the two-bottle choice procedure, adolescents consumed significantly more ethanol than adults. Fo r total ethanol consumption (g/Kg of EtOH), ANOVA revealed significant differences for the ontogenic period factor [F(1,57) = 78.77; *p* < 0.0001] and significant interaction between factors [F(8,456) = 4.707; *p* < 0.0001], with *post-hoc* differences observed in sessions S1, S2, and S7 (*p* < 0.0001; Fig. 1E). During quinine-adulterated ethanol access (g/Kg of EtOH), ANOVA revealed that there was a difference for the ontogenic period factor [F(1,57) = 29.83; *p* < 0.0001], with no significant interaction between the factors [F(3,171) = 1.088; *p* > 0.05], adolescents consumed more ethanol across all quinine sessions (Fig. 1F). These findings suggest that ethanol intake persists longer in adolescent mice despite aversive adulteration.

Analyses of ethanol preference similarly revealed heightened ethanol motivation in adolescents. For ethanol preference in EtOH periods 1 (S1-S6) and 2 (S7-S9), ANOVA revealed a significant difference for the ontogenic period factor [F(1,57) = 33.82; *p* < 0.0001], with no significant interaction between the factors [F(8,456) = 0.8097; *p* < 0.0001] (Fig. 1G). During the quinine phase (SQ1–SQ3), ANOVA again revealed difference for the ontogenic period factor [F(1,57) = 56.93; *p* < 0.0001] and a significant interaction between factors [F(3,171) = 4.25; *p* < 0.01]. *Post-hoc* analysis showed differences for all sessions of the quinine period (*p* < 0.001), with adolescents displaying significantly higher preference in all quinine sessions (Fig. 1H).

As described above, prior to performing the principal component analysis (PCA), a Pearson correlation matrix was constructed using all OCL and two-bottle choice variables (Fig. I). Variables exhibiting high correlations (|*r*| > 0.8) were excluded to reduce redundancy. The PCA was then conducted using the remaining variables to further characterize age-dependent behavioral profiles. Twelve components were generated; five had eigenvalues greater than 1, and together they accounted for 73.1% of the total variance. CP1 (17.6%, eigenvalue = 3.13), primarily driven by early ethanol and quinine-ethanol consumption variables (S1 and SQ2), differed significantly between adolescents and adults (*p* < 0.0001; Fig. 1J). CP2 (17.1%, eigenvalue = 1.66), loaded by OCL-nose pokes and reinforcement parameters, also differed between age groups (*p* < 0.05; Fig. 1K). CP3 (12.7%) and CP4 (12.0%) accounted for later quinine session variance and showed no age-related effects. CP5 (8.5%, eigenvalue = 1.03), associated with early two-bottle choice sessions, exhibited significant adolescent-adult divergence (*p* < 0.001; Fig. S1). Across components, adolescents showed stronger covariance among impulsivity measures and ethanol intake, even under aversive conditions.

Finally, Mann–Whitney test performed on Fos immunohistochemistry data obtained after the final ethanol session (S9) revealed no age-related differences in total Fos-positive neuronal counts in the insular cortex (*p* > 0.5) (Supplementary Figs. S2–S3).

### 4.2 Deficit in inhibitory control in adolescent mice is associated with altered insular cortex activation

To investigate the neural correlates underlying age-dependent impulsivity and ethanol-related behaviors, we quantified neuronal activation in the anterior insular cortex following completion of the OCL task (Fig. 2B–D). Fos immunoreactivity was assessed to evaluate overall neuronal activation (NeuN+/Fos+) (Fig. 2E) as well as activation within parvalbumin-positive (PV+) GABAergic interneurons

Adolescent mice exhibited significantly greater Fos expression in the insular cortex compared with adults (*p* < 0.001; Fig. 2F). Consistent with this, the number of double-labeled NeuN+/Fos+ neurons was also higher in adolescents (*p* < 0.01 Fig. 2H), resulting in a greater percentage of activated neurons overall (*p* < 0.05; Fig. 2I).

To evaluate PV interneuron activation, we quantified the proportion of PV+ cells co-expressing Fos (PV+/Fos+) (Fig. 2J). Only a small fraction of PV interneurons was activated in both age groups (Fig. 2M). Adolescents showed a lower percentage of activated PV interneurons (1.87 ± 0.14) relative to adults (2.92 ± 0.62), despite exhibiting a higher total number of Fos-labeled cells overall (*p* < 0.05). The total number of PV+ interneurons did not differ between groups (*p* > 0.05; Fig. 2L). In contrast, adults displayed a significantly greater number of PV+/Fos+ neurons (*p* < 0.01) (Fig. 2K) and a higher percentage of PV interneuron activation (*p* < 0.05) compared with adolescents (Fig. 2N).

Together, these findings suggest that while adolescents exhibit increased global neuronal activation within the insular cortex, adults display a preferential engagement of PV interneurons. This divergence suggests developmental differences in inhibitory circuit recruitment that may contribute to heightened impulsivity and ethanol-related behaviors during adolescence.

### 4.3 Activation of parvalbumin-positive interneurons in the insular cortex enhances inhibitory control

To directly assess the causal contribution of PV+ GABAergic interneurons in the insular cortex to impulsivity-related behaviors, we expressed Cre-dependent DREADDs (Gq- or Gi-coupled human muscarinic receptors) in PV-Cre transgenic mice. We evaluated their performance in the OCL task. Thirty minutes before testing, mice received an intraperitoneal injection of CNO (1 mg/kg), a dose shown to effectively activate both Gq- and Gi-DREADDs.

During OCL training, two-way ANOVA on active nose-pokes (ANP) indicated a significant difference for the sessions factor [F(8,96) = 20.99; *p* < 0.0001] and significant interaction between factors [F(16,96) = 1.833; *p* = 0.0374]. *Post-hoc* analysis indicated no differences between groups across sessions (*p* > 0.05) (Fig. 3B). Similarly, ANOVA indicated differences for the DREADD factor [F(12,96) = 3.383; *p* < 0.001] and a significant interaction between the factors [F(16,96) = 6.191; *p* < 0.0001]. The *post-hoc* analysis showed no differences between the groups throughout the sessions, for the number of rewards and nose-pokes (*p* > 0.05) (Fig. 3C).

In the OCL test session, however, chemogenetic manipulation produced robust behavioral effects. ANOVA revealed significant group differences in both ANP [F(2,12) = 13.60; *p* < 0.0001] and REW [F(2,12) = 8.82; *p* < 0.0001]. *Post-hoc* comparisons showed that Gi-DREADD inhibition of PV interneurons increased impulsive responding (higher ANP) relative to both Control (*p* < 0.05) and Gq groups (p < 0.001), whereas Gq-DREADD activation reduced impulsive responses (Fig. 3D). Reward outcomes followed a similar pattern, with Gq activation decreasing reinforcement acquisition relative to both Control and Gi groups (Fig. 3E).

These findings demonstrate that the activity of insular PV+ interneurons exerts bidirectional control over impulsive responding. Enhancing PV interneuron activity (Gq-DREADDs) improves inhibitory control by suppressing inappropriate nose-pokes, whereas inhibiting these interneurons (Gi-DREADDs) diminishes inhibitory control and promotes impulsive behavior. Overall, PV interneurons in the insular cortex play a critical modulatory role in regulating impulse-related actions during the OCL task.

### 4.4 Chemogenetic modulation of PV+ interneurons in the insular cortex alters ethanol-seeking responses

To determine whether insular PV+ GABAergic interneurons regulate ethanol seeking behavior, PV-Cre mice injected with AAV-EF1a-DIO-hM3Dq or control vectors (n = 8 per group) were subjected to a forced-ethanol consumption protocol (12 h / 6 h / 4 h / 2 h). Behavioral testing was conducted 30 minutes following CNO administration (1 mg/kg; Fig. 4A).

During OCL training, two-way ANOVA on active nose-pokes (ANP) indicated a significant difference for the sessions factor [F(8,160) = 63.83; *p* < 0.001] and significant interaction between factors [F(16,160) = 3.483; *p* < 0.05]. *Post-hoc* analysis indicated differences between the Control and Gq groups for the FR5-01, 02 and 03 sessions (*p* < 0.05) (Fig. 4B). For rewards (REW), ANOVA revealed differences for the DREADD factor [F(8,160) = 13.11; *p* < 0.05] and significant interaction between the factors [F(16,160) = 2.046; *p* < 0.001]. However, the *post-hoc* analysis did not demonstrate differences between the groups, throughout the sessions, for the number of rewards (*p* > 0.05) (Fig. 4C).

In the OCL test, ANOVA indicated significant group effects for ANP [F(2,21) = 3.452; *p* < 0.05], with *post-hoc* analysis showing increased impulsive responding in Gi relative to Gq mice (*p* < 0.05; Fig. 4D). For REW, ANOVA also revealed significant group differences [F(2,21) = 15.59; *p* < 0.0001], with Control vs. Gi (*p* < 0.01) and Gi vs. Gq (*p* < 0.0001) pairs differing significantly (Fig. 4E).

During the final forced-ethanol exposure session (T = 2 h), ethanol consumption (g / kg EtOH) did not differ between groups [F(2,20) = 2.586; *p* > 0.05; Fig. 4H]. However, during the ethanol self-administration test, ANOVA revealed significant group differences for both ANP [F(2,20) = 4.905; *p* < 0.05] and REW [F(2,20) = 4.286; *p* = 0.0282]. *Post-hoc* analysis revealed significant differences between Gi and Gq for both outcomes (p ≈ 0.02–0.03; Fig. 4F–G).

To rule out nonspecific effects of CNO on locomotor activity, we performed an open-field assessment. No significant group differences were observed in total distance traveled [F(2,20) = 0.6351; *p* > 0.5] or average speed [F(2,20) = 0.2183; *p* > 0.5] (Supplementary Fig. S4).

To evaluate PV interneuron activation, we quantified the proportion of PV+ cells co-expressing Fos (PV+/Fos+). ANOVA revealed significant group differences [F(2,12) = 11.47; *p* < 0.01]. *Post-hoc* comparisons showed significant differences between th Gi *vs.* Gq groups (*p* < 0.01) and Control *vs.* Gi (*p* < 0.01). No significant difference was observed between Control *vs*. Gq (*p* > 0.05) for the percentage of activated PV interneurons (Fig. 4I).

These results demonstrate that chemogenetic modulation of insular PV interneurons selectively alters ethanol-seeking behavior without affecting locomotion. Specifically, activation of PV+ interneurons (Gq-DREADDs) reduces ethanol-seeking responses. In contrast, their inhibition (Gi-DREADDs) increases operant responding for ethanol, underscoring their essential role in regulating motivated and reward-driven behaviors.

## 5. Discussion

Adolescence is a critical developmental window marked by heightened impulsivity and vulnerability to substance use, yet the neural mechanisms underlying these behaviors remain partly understood. In this study, we provide a comprehensive analysis of how the insular cortex, specifically its subpopulation of PV+ GABAergic interneurons, modulates impulsivity and ethanol-seeking behaviors across developmental stages in mice [37]. Consistent with previous studies[38]–[40], our results show that adolescent mice displayed higher rates of impulsivity compared to adulthood, as evidenced by increased active nose-pokes on the OCL test. In addition, adolescents also exhibit significantly greater ethanol consumption compared to adults with higher ethanol preference, even in the presence of aversive stimuli such as quinine on the two-bottle choice protocol. At the neural level, we observed that adolescent mice display elevated overall neuronal activation in the insular cortex, as indicated by increased Fos expression and a higher proportion of double-labeled NeuN+/Fos+ cells. However, activation of PV+ interneurons was lower in adolescents compared to adults, suggesting an imbalance towards higher activation in the insular cortex, probably related to a developmental lag in the maturation of inhibitory microcircuits. Critically, we found that chemogenetic inhibition of PV+ interneurons in the insular cortex of adult mice reinstated a lack of inhibitory control, increasing both impulsivity and ethanol-seeking behaviors, whereas their activation produced the opposite effect. Our findings provide a neurobiological mechanism for the heightened impulsivity and increased vulnerability to ethanol consumption.

In our study, we first validated the differences in impulsive behaviors between adult and adolescent animals in the OCL. We observed larger numbers of target-driven behaviors, such as active nose pokes, which may reflect an inability to control impulsive behavior in adolescent animals [41]–[43]. In adults, a decrease in the frequency of actions such as bar pressing or nose-pokes following the initial OCL session indicates the development of learning and adaptive behavior [44]. Considering that impulsivity is a predictor of drug use [15], [45], [46], we hypothesize that adolescents would be more susceptible to the development of alcohol-related disorders. The persistence of impulsive behaviors and elevated consumption in adolescents, despite aversive stimulus (quinine), observed in our study, is in line with prior evidence indicating high rates of impulsive choices and high alcohol consumption in adolescents compared to [42], [47], [48] adults [49]. The transition between adolescence and adulthood converges to behaviors with lower impulsivity scores, with a decrease in consumption and problematic use of alcohol and other drugs [50]. However, the basal inhibitory function typical of adolescence is strongly impaired by alcohol use at this stage, increasing the vulnerability of developing alcohol and other drug use disorders in adulthood [51]–[53]. Interestingly, rats with a higher preference and ethanol consumption, when compared to animals with low consumption and preference, have greater sensitivity to delayed rewards, that is, an undervaluation of late reinforcements [54], which could help explain the results of our OCL and 2BC experiments.

Notwithstanding that impulsivity can increase drug consumption, prior drug exposure may in turn amplify impulsive-related behaviors. In rats treated with d-methamphetamine and subjected to an omission and delayed discount protocol, the number of omissions remained unchanged, while active lever presses increased across subsequent sessions [45]. These findings indicate a bidirectional relationship, whereby impulsivity promotes drug intake and drug use further intensifies impulsive-related behavior.

Functional alterations in the insular cortex have been closely linked to impulsivity and impaired decision-making under risk-taking conditions [54]–[57]. Activation of the insular cortex has been shown to enhance the motivation for alcohol consumption, influencing the severity of drinking [58]. Furthermore, chemogenetic manipulation of insular cortex glutamatergic neurons modulated alcohol consumption [32], [59], impulsive behaviors, and risk choice[60]. Our findings extend this knowledge by highlighting the contribution of PV interneurons in the insular cortex to impulsivity-related behavior and alcohol consumption. Specifically, we observed an increase in the number of Fos+ cells in the insula of adolescent animals, suggesting a correlation between the activation of the insular cortex and the emergence of impulsive-related behaviors. In addition, we found less activated PV in the insular cortex of adolescent mice than in adults, suggesting a lack of local inhibition.

To assess the role of PV+ interneurons in impulsivity, we used chemogenetics to modulate their activity in adult animals. Inhibition of PV+ interneurons increased impulsive behaviors, while activation of these interneurons decreased them, demonstrating a causal role of PV+ interneuron activity in the insular cortex in impulsive-related behavior. These results align with previous studies, which have shown that PV+ interneurons regulate inhibitory control in the Five-Choice test [61] The implication of PV+ interneurons suggests late insular maturation and neuronal hyperexcitability of this region in adolescent animals [51]–[53], [62]. Immature PV+ interneurons may promote an increased subjective valence of reward and inefficient interoceptive processing [29], [57], [63], [64]. Ultimately, inhibitory microcircuitry dysfunctions, involving PV+ interneurons, may be able to increase the activity of pyramidal neurons and promote cognitive-behavioral impairments [27], [65].

Interestingly, when neuronal activation was assessed after ethanol consumption, the previously observed age-related differences in insular activation were no longer detectable. This could be due to alcohol-induced increases in insular activation that mask developmental differences, or to the duration of the 4-hour consumption session that may have masked the percentage of neuronal activation. Our findings also showed that adolescent animals are less sensitive to taste aversion than adult animals[50], [66], [67], possibly due to ongoing maturation of insular circuits involved in taste and aversion processing [67]–[69]. The neurofunctional organization of the insula during adolescence is a complex and dynamic process, involving age-dependent changes across different subregions and neural circuits [50], [66], [70], which could explain these differences in aversive sensitivity. The insular cortex plays a central role in taste-visceral integration, perception of aversive flavors, and conditioned taste aversion[69], [71], [72]. Decreasing the activity of the insular cortex diminished the malaise caused by the acute administration of lithium chloride [73], suggesting that the insula plays a crucial role in the perception and processing of aversive visceral sensations [74], [75].

Overall, our findings underscore the importance of the insular cortex and its PV interneurons in regulating impulsive and ethanol-related behaviors across development. The maturation of inhibitory circuits, including PV-mediated control over pyramidal neuron activity, appears central to the transition from adolescent vulnerability to adult behavioral stability. These data highlight PV interneurons as potential targets for interventions aimed at mitigating impulsivity and reducing vulnerability to ethanol misuse.

## 6. Conclusion

Our findings identify the insular cortex as a critical regulator of impulsivity and ethanol-related behaviors, highlighting the essential contribution of insular parvalbumin-positive (PV+) interneurons to developmental vulnerability for alcohol use disorder (AUD). Given the insula’s central role in integrating interoceptive, motivational, and aversive information, disruptions in PV-mediated inhibitory control may represent a key mechanism contributing to adolescent vulnerability to AUD. Future studies should examine the longitudinal developmental trajectories of insular GABAergic function, assess subregion-specific circuit interactions, and explore how PV-mediated inhibition shapes the maturation of reward and aversion pathways. Elucidating these mechanisms may enable the development of targeted strategies aimed at enhancing inhibitory control and reducing addiction risk during adolescence.

## Data Availability Statement

The datasets generated and/or analyzed during the current study are available from the corresponding author upon reasonable request. All data supporting the findings of this study are included in the article and its supplementary materials.

## Ethics Approval Statement

All experimental procedures involving animals were conducted in accordance with the guidelines of the Principles of Care of Laboratory Animals. Protocols were approved by the Ethics Committee on Animal Use of the Federal University of São Paulo (CEUA #9191091219) and by the local Animal Experimentation Ethics Committee C2EA-05 Charles Darwin of Sorbonne Université, under the European Council Directive 2010/63/EU, as authorized by the French Ministry of Higher Education and Research.

## Competing Interests

The authors declare that they have no competing interests.

## Author Contributions (CRediT taxonomy)

Fernando B. Bezerra: Conceptualization; Methodology; Investigation; Formal Analysis; Visualization; Writing – Original Draft.

G.J.D. Fernandes: Investigation; Writing – Review & Editing

A. Anjos-Santos: Investigation; Writing – Review & Editing

C.A. Favoretto: Investigation; Writing – Review & Editing

B.T. Rodolpho: Writing – Review & Editing

P. Palombo: Visualization, Writing – Review & Editing

V. Vialou: Resources; Supervision; Writing – Review & Editing.

K.P. Abrahao: Conceptualization; Supervision; Review & Editing.

F.C. Cruz: Conceptualization; Funding Acquisition; Project Administration; Supervision; Writing – Review & Editing.

## Acknowledgments

The authors thank the technical staff of the Laboratory of Molecular and Behavioral Neuroscience at UNIFESP for their essential support. We also thank the Center for Development of Experimental Models in Biology and Medicine (CEDEME) for providing animal facilities and technical assistance. This work was supported by funding from the São Paulo Research Foundation (FAPESP 2019/24073-3; 2018/15505-4; 2019/01686-0), the Brazilian National Council for Scientific and Technological Development (CNPq 314336/2023-0), and the Coordination for the Improvement of Higher Education Personnel (CAPES). We are grateful to the imaging facilities teams at Sorbonne Université for their assistance with histological analyses.

## Notes

### Competing Interest Statement

The authors have declared no competing interest.

### Summary of Updates

The only modification in this revision was a correction to the title. The title has been changed to: The Role of Anterior Insular Parvalbumin GABAergic Interneurons in Impulsivity-Related and Alcohol-Seeking Behaviors. The previous title (The Role of Insular Anterior Parvalbumin...), contained an incorrect word order. No changes were made to the manuscript content.

