## Supplementary material for "The Role of Anterior Insular Parvalbumin GABAergic Interneurons in Impulsivity-Related and Alcohol-Seeking Behaviors": Supp Fig Captions

**Supplementary Figure Captions**

**Figure S1**. Principal Component Analysis. Principal Component 3 (PC3) was loaded by consumption in the 3rd session of EtOH + Quinine (g/Kg/EtOH – SQ3) and by preference in this same session (%PREF – SQ3). Principal Component 4 (CP4) was formed by the consumption of ethanol in the 1st session of EtOH + Quinine (g/Kg/EtOH – SQ1). by preference in this same session (%PREF – SQ1) and by preference in the first ethanol session (%PREF – S1 Principal Component 5 (CP5) was formed by the consumption in the 2nd session of EtOH (g/Kg/EtOH – S2), by the preference in this same session (%PREF – S2) and also by the consumption of ethanol in the 1st session of the Two-Bottle Choice (g/Kg/EtOH – S1). *** *p* < 0.001. Adolescents, *n* = 8; Adults, *n* = 8. Data shown as means ± standard deviation of the mean (SEM).

**Figure S2**. Neuronal activation (number of Fos-positive cells/mm^2^) during the last session of the two-bottle free choice of ethanol intake (Adolescent mice in blue; Adult mice in red). **p* > 0.05. Adolescents, *n* = 8; Adults, *n* = 8. Data shown as means ± standard deviation of the mean (SEM).

**Figure S3**. Representative images of Fos-positive cells in the insular cortex. In A, illustration of a brain coronal slice, highlighting the insular cortex (in red) (adapted from Paxinos & Watson, 2007). In B, photomicrograph of the insular cortex of an adolescent mouse. In C, photomicrograph of the insular cortex of an adult mouse.

**Figure S4**. Locomotor activity in the open field. In A, total distance traveled (m) during the open field test. In B, average speed (m/s) of mice during the open field test. [control DREADD in black; inhibitory DREADD (Gi) in red; excitatory DREADD (Gq), in blue]. **p* < 0.05; *n* = 4 mice per group. Data shown as means ± SEM.
