## Supplementary material for "The Role of Anterior Insular Parvalbumin GABAergic Interneurons in Impulsivity-Related and Alcohol-Seeking Behaviors": Supp Figures

### Supplementary Results

#### Experiment 1: Figure S1

**PCA**

**A**

**PC3**

**1.0**

**Correlation coefficients**

**0.8**

**0.6**

**0.4**

**0.2**

**0.0**

**PC3**

**g/Kg/EtOH - SQ3	%PREF - SQ3**

**B**

**4**

**2**

**Factorial load**

**0**

**-2**

**-4**

**Adolescents	Adults**

**A	PC4	B**

**PC4**

**1.0	4**

**Correlation coefficients**

**0.5	2**

**Factorial load**

**0.0	0**

**-0.5	-2**

**-1.0**

**g/Kg/EtOH - SQ1 %PREF - SQ1	%PREF - S1**

**-4**

**Adolescents	Adults**

**A**

**PC5**

**1.0**

**Correlation coefficients**

**0.8**

**0.6**

**0.4**

**0.2**

**0.0**

**PC5**

**g/Kg/EtOH - S2	%PREF - S2**

**B**

**6**

✱✱✱

**4**

**Factorial load**

**2**

**0**

**-2**

**Adolescents	Adults**

#### Experiment 2: Figure S2

**300**

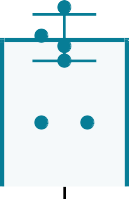

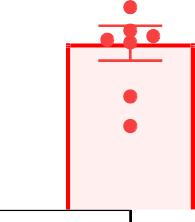

**Fos^+^ / mm^2^**

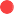

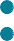

**200**

**100**

**0**

**Adolescents	Adults**

**Experiment 2: Figure S3**

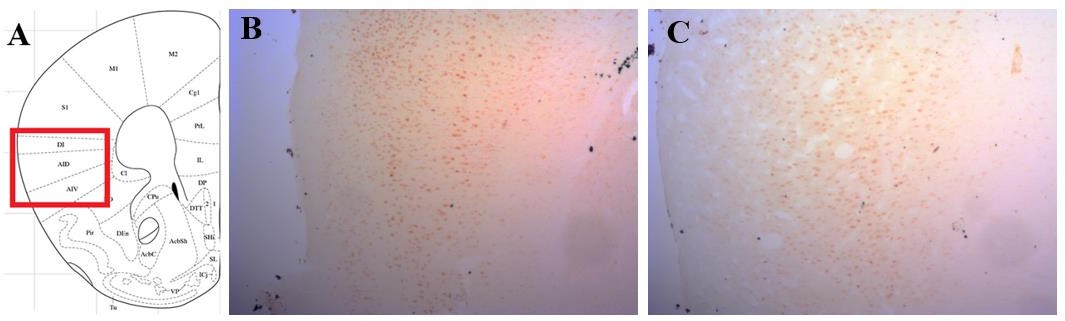

#### Experiment 4: S4

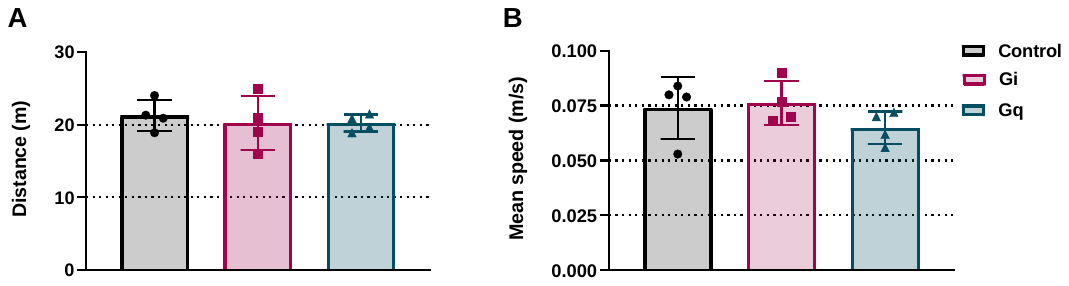
